# DrImpute: Imputing dropout events in single cell RNA sequencing data

**DOI:** 10.1101/181479

**Authors:** Il-Youp Kwak, Wuming Gong, Naoko Koyano-Nakagawa, Daniel J. Garry

## Abstract

The single cell RNA sequencing (scRNA-seq) technique began a new era by allowing the observation of gene expression at the single cell level. However, there is also a large amount of technical and biological noise. Because of the low number of RNA transcriptomes and the stochastic nature of the gene expression pattern, there is a high chance of missing nonzero entries as zero, which are called dropout events. However, many statistical methods used for analyzing scRNA-seq data in cell type identification, visualization, and lineage reconstruction do not model for dropout events. We have developed DrImpute to impute dropout events, and it improves many of the statistical tools used for scRNA-seq analysis that do not account for dropout events. Our numerical studies with real data demonstrate the promising performance of the proposed method, which has been implemented in R.

## Background

DNA sequencing technology is growing rapidly, and next generation sequencing approaches for high-throughput RNA sequencing are phenomenal. Bulk RNA sequencing (bulk RNA-seq) technology performs high throughput sequencing using bulk populations of millions of input cells, which implies that the resulting expression value for each gene is the average expression value of a large population of input cells ([1], [2]). Thus, bulk RNA-seq is suitable for revealing a global view of averaged gene expression levels. However, the bulk RNA-seq method is not capable of quantifying the RNA contents of very few cells and yields biased results when mixed cell types exist in the samples ([2]). For example, bulk RNA-seq is unable to accurately reveal the transcriptome of the cells from the early embryonic developmental stage where there exist multiple lineages with relatively small numbers of cells. Recently, scRNA-seq was developed to enable a wide variety of transcriptomic analyses at the single cell level ([3], [4], [5], [6], [7], [8]). The major areas in scRNA-seq research include characterizing the global expression profiles of rare cell types, discovering novel cell populations, and reconstructing cellular developmental trajectories ([9], [10], [6], [8], [11], [12]). Accordingly, many statistical methods have been developed for clustering cell populations, visualizing cell-wise hierarchical relationships, and predicting lineage trajectories ([13], [11], [14], [15], [16], [17], [18], [19], [20], [21], [22]).

However, scRNA-seq has a relatively higher noise level than bulk RNA-seq especially due to so-called dropout events ([2], [23], [24]). Dropout events are special types of missing values (a missing value is an instance wherein no data is present for the variable), caused both by low RNA input in the sequencing experiments and by the stochastic nature of the gene expression pattern at the single cell level ([25], [26], [23], [27]). Consequently, these dropout events become zeros in the resulting gene-cell expression matrix and they mix with the true zeros where a gene is not expressed in a cell at all. However, most statistical tools developed for scRNA-seq analysis do not explicitly address these dropout events ([13], [11], [15], [16], [28], [19], [20], [21], [22]). We hypothesize that imputing the missing expression values caused by the dropout events will improve the performance of cell clustering, data visualization, and lineage reconstruction.

The gene expression data from bulk RNA-seq (or microarrays) also suffer from a missing value problem ([29]). Various statistical methods have been proposed to estimate the missing values in the data ([29], [30], [31], [32], [33]). These missing value imputation methods can be categorized as five general strategies, as follows: 1) *mean imputation* estimates missing entries by simply averaging gene-level or cell-level expression levels ([29], [30]); 2) *hot deck imputation* predicts missing values from similar entries using a similarity metric among genes (KNNImpute, [31]); 3) *model based imputation* employs statistical modeling to estimate missing values (GMCimpute, [30]); 4) *multiple imputation* methods predict missing entries multiple times and combine the results to produce final imputation (SEQimpute, [32]); and 5) *cold deck imputation* uses side information such as gene ontology to facilitate the imputation process (GOkNN, GOLLS, [33]).

However, the imputation methods developed for bulk RNA-seq data may not be directly applicable to scRNA-seq data. First, much larger cell-level variability exists in scRNA-seq, because scRNA-seq has cell-level records for gene expression; on the other hand, bulk RNA-seq data have the averaged gene expression of the bulk population of cells. Second, dropout events in scRNA-seq are not exactly missing values; dropout events have zero expression, and they are mixed with real zeros. In addition, the proportion of missing values in bulk RNA-seq data is much smaller. Therefore, a dropout imputation model for scRNA-seq is needed.

There are three previous studies for imputing dropout events ([34], [35], [36], [37]). BISCUIT iteratively normalizes, imputes, and clusters cells using the Dirichlet process mixture model ([35], [36]). Zhu et al. proposed a unified statistical framework for both single cell and bulk RNA-seq data ([34]). The scImpute infers dropout events based on dropout probability, using weighted LASSO with genes similar to those of impute dropout events. However, none of these three methods demonstrate how imputing dropout events could possibly improve the current statistical methods that do not account for dropout events.

In the present study, we designed a simple, fast hot deck imputation approach, called DrImpute, for estimating dropout events in scRNA-seq data. DrImpute first identifies similar cells based on clustering, and imputation is performed by averaging the expression values from similar cells. To achieve robust estimations, the imputation is performed multiple times using different cell clustering results followed by averaging multiple estimations for final imputation. We demonstrated on seven published scRNA-seq datasets that imputing the dropout entries significantly improved the performance of existing tools (pcaReduce, SC3, t-SNE, PCA, Monocle, and TSCAN) in regard to cell clustering, visualization, and lineage reconstruction. Moreover, DrImpute also performed better than Clustering through Imputation and Dimensionality Reduction (CIDR), Zero Inflated Factor Analysis (ZIFA), and scImpute in accounting for dropout events in scRNA-seq data ([14], [18]).

## Results

### Cell clustering

Discovering distinct cell types from a heterogeneous cell population (cell clustering) is one of the most important applications of scRNA-seq ([2]). Several methods, such as pcaReduce, SC3, and t-SNE followed by k-means (t-SNE/kms), have been developed for clustering scRNA-seq data ([13], [11], [38]). However, these methods do not explicitly address the dropout events or the missing values existing in the scRNA-seq data. We hypothesized that 1) preprocessing the scRNA-seq data by imputing the dropout events via DrImpute will improve the accuracy of these clustering methods and 2) the performance of the existing tools combined with DrImpute will perform better than with CIDR or scImpute in addressing dropout events ([14], [37]).

First, we evaluated whether imputing the dropout events using DrImpute before applying pcaReduce, SC3, and t-SNE would improve the accuracy of cell type identification. We compared the clustering performance of these methods with and without imputing dropout events by DrImpute, on five published scRNA-seq datasets ([10], [6], [39], [7], [40]). Using the cell types reported in the original publications as the ground truth and the Adjusted Rand Index (ARI) as the performance metric, we found that preprocessing the scRNA-seq datasets with DrImpute significantly improved the clustering performance of pcaReduce with the M and S options (pcaR_M and pcaR_S) on all five tested datasets; improved the performance of t-SNE followed by k-means (t-SNE/kms) on four datasets; and improved the performance of SC3 on three datasets (Figure 1a). Second, we also found that combining DrImpute with t-SNE/kms showed significantly better clustering performance than CIDR on three of five datasets and on scImpute followed by t-SNE/kms on four of five datasets (Figure 1a).

**Figure 1.**
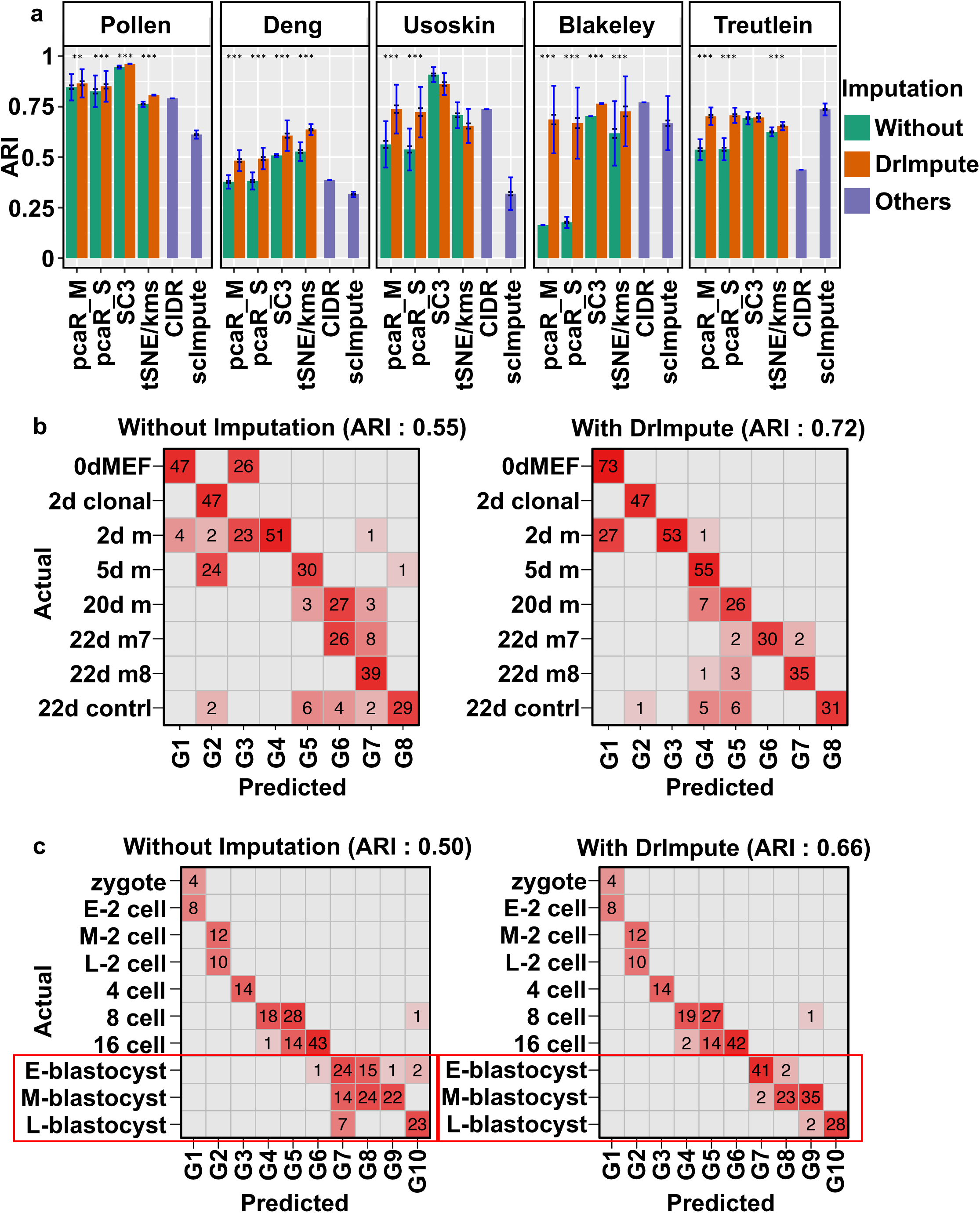
DrImpute significantly improved the performance of the existing tools for cell type identification. **(a)** The average adjusted Rand index (ARI) of 100 repeated runs of pcaR_M (pcaReduce with the M option), pcaR_S (pcaReduce with the S option), SC3, t-SNE/kms (t-SNE followed by k-means), CIDR and scImpute, on five scRNA-seq datasets. Blue interval represents one plus or minus standard deviation of the category. Black interval represents one plus or minus standard error of the category. Wilcoxon rank sum test was utilized to compare the ARIs from different tools (⁎⁎: 0.01 ≤ p value < 0.001, ⁎⁎⁎ p value < 0.001). **(b-c)** The confusion matrix for **(b)** iN reprograming ([40]) using pcaReduce (option S) and **(c)** mouse preimplantation embryo ([6]) using t-SNE followed by k-means. Y axis represents ground truth cluster groups reported in the original study and X axis represents predicted groups. Left and right panels, respectively, represent the confusion matrix according to the clustering results without and with preprocessing the scRNA-seq data using DrImpute. The ARI was computed between the original and predicted cell groups.

Figure 1b shows a confusion matrix of the ground truth cell labels and cell clusters predicted by pcaReduce (option S) on the scRNA-seq dataset of induced neuronal (iN) reprograming, without (left panel) and with (right panel) imputing the dropout events using DrImpute ([40]). We observed a clearer diagonal pattern in the confusion matrix with the imputation, as supported by an improvement of ARI, from 0.55 to 0.72. As another example, Figure 1c shows the confusion matrix of the ground truth labels and cell clusters predicted by t-SNE/kms on a dataset of mouse preimplantation embryos ([6]). We found that imputing the dropout events helped t-SNE/kms more accurately cluster the cells from the blastocyst stages, as evidenced by an increase in mean ARI from 0.50 to 0.66.

We further assessed whether preprocessing scRNA-seq by imputing dropout events would produce more consistent clustering results. We evaluated the robustness of the clustering results with and without imputing the dropout events using DrImpute. We hypothesized that preprocessing the scRNA-seq with DrImpute would help the clustering methods detect more robust and consistent subpopulations. For each dataset, we randomly sampled 100 genes (gene level down-sampling), or one-third of the total cells (cell level down-sampling), and we clustered the cells using each of the clustering methods with and without preprocessing the down-sampled dataset using DrImpute. This process was repeated 100 times, and we compared how consistent the clustering results were after down-sampling the genes or cells as measured by cross ARI (see Methods). For both the gene and cell down-sampling experiments, we found that preprocessing of the scRNA-seq datasets with DrImpute significantly improved the robustness of the cell type identification of SC3, t-SNE/kms, and pcaR_M / pcaR_S on 80% of the tested cases (Supplementary Figure 3a and 3b).

In summary, these results suggested that in 50 out of 60 tested cases, preprocessing the scRNA-seq datasets by imputing the dropout events using DrImpute significantly improved the accuracy or the robustness of clustering methods that did not specifically address dropout events. Combined with DrImpute, pcaReduce, SC3, and t-SNE followed by k-means also had significantly better performance than CIDR in terms of both the accuracy (13 out of 20 cases) and the robustness of cell clustering.

### Cell visualization

Principal component analysis (PCA) and t-SNE are the two most popular methods for visualizing scRNA-seq in a two-(2D) or three-dimensional (3D) space ([28]). However, neither PCA nor t-SNE explicitly addresses dropout events. Recently, Zero Inflated Factor Analysis (ZIFA) was developed to factorize and visualize scRNA-seq data while also addressing dropout events. We hypothesized that with the preprocessing of scRNA-seq data by imputing the dropout events using DrImpute, the generic dimension reduction methods (PCA and t-SNE) would generate better factorization/visualization results than without imputation.

To evaluate the accuracy of the dimension reduction on a 2D space, we first estimated how discriminatively the cells from one population (using the class label reported in the original publication) separated from other populations on the 2D space. For each dimension reduction result, we used the 2D coordinates of 90% of cells as the feature to train a linear support vector machine classifier, and we predicted the class label for the remaining 10% of the cells. The above process was repeated ten times, and the overall prediction accuracy (10-fold cross validation accuracy) was used to quantitatively measure the separability of different populations on a 2D space.

We compared the performance of PCA and t-SNE with and without DrImpute preprocessing as well as ZIFA and t-SNE with scImpute on five published scRNA-seq datasets. We observed significant improvements in both PCA and t-SNE with DrImpute (Figure 2a). Moreover, on three datasets where ZIFA had better separation than PCA, preprocessing the data with imputation employing DrImpute helped PCA achieve significantly better performance than ZIFA in separating the cell populations (Figure 2a).

**Figure 2.**
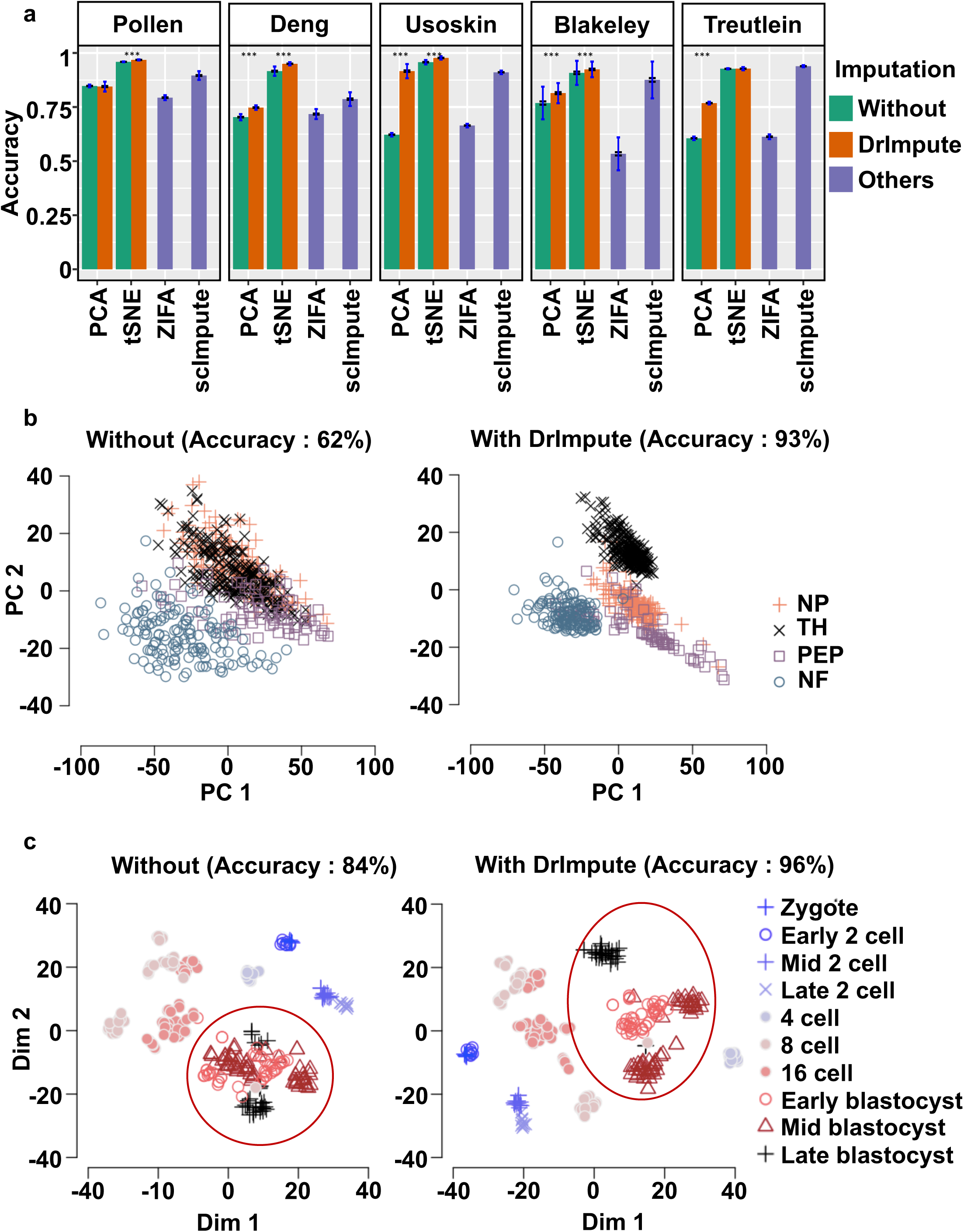
DrImpute significantly improved the performance of PCA and t-SNE in visualizing scRNA-seq data. **(a)** The barplots of average accuracy of separating the cell subpopulations on the 2D space. Blue interval represents one plus or minus standard deviation of the category. Black interval represents one plus or minus standard error of the category. Wilcoxon rank sum test was performed to compare the average ARIs from different tools. Eight out of 10 cases are statistically significant (⁎⁎⁎ *p* value < 0.001). **(b)** Visualization of four groups of mouse neural single cells (NP, TH, PEP, and NF) ([39]) using PCA. Left and right panels, respectively, show the 2D visualization of single cells without and with preprocessing the scRNA-seq data using DrImpute. **(c)** Visualization of mouse preimplantation embryo ([6]) using t-SNE. Left and right panels, respectively, show the 2D visualization of single cells without and with preprocessing the scRNA-seq data using DrImpute. The classification accuracy was computed by using the 2D coordinates of each dimension reduction results.

Figure 2b shows the cell expression profiles of four types of mice neurons (non-peptidergic nociceptors [NP], tyrosine hydroxylase containing [TH], peptidergic nociceptors [PEP], and neurofilament containing [NF]) using PCA without (left) and with (right) imputing the dropout events using DrImpute ([39]). Without imputing the dropout events with DrImpute, the NP, TH, and PEP groups were visually indistinguishable in the 2D space. However, after applying DrImpute, all four groups were visually separated, as demonstrated by an accuracy increase from 62% to 93%. Figure 2c shows the cell expression profiles of mouse preimplantation embryos using t-SNE ([6]). As seen in the red circled area, the stages of early, mid, and late blastocyst were more clearly distinguished after preprocessing the scRNA-seq data with DrImpute, as supported by an accuracy increase from 84% to 96%.

In summary, we found that preprocessing the scRNA-seq datasets by imputing the dropout events using DrImpute significantly improved the accuracy of visualization, as shown from the significant results on 20 of 30 cases in our study. The generic dimension reduction methods (PCA and t-SNE) on imputed datasets using DrImpute also performed significantly better than ZIFA, which was specifically designed for scRNA-seq data considering dropout events.

### Lineage reconstruction

The third common task for single cell RNA-seq analysis is to reconstruct the lineage trajectories and infer the differentiated and progenitor states of the single cells. For example, Monocle and TSCAN were designed to infer pseudotime from the biological cellular process ([19], [20]). However, neither method accounts for dropout events. We hypothesized that inferring the pseudotime on scRNA-seq data preprocessed using DrImpute could improve the accuracy of pseudotime ordering.

We compared the performance of pseudotime inference with and without imputing the dropout events on three published time series scRNA-seq datasets, mouse preimplantation embryonic development data, human preimplantation embryonic development data, and mouse early mesodermal development data ([6], [41], [42]). We used the reported time labels as the ground truth and evaluated the performance of pseudotime inference by comparing the time labels and pseudotime. The consistency between the time labels and pseudotime ordering was measured by the Pseudo-temporal Ordering Score (POS, [20]) and Kendall’s rank correlation score.

We found that both Monocle and TSCAN had significantly better performance on pseudotime inference on all three tested datasets if the scRNA-seq data were preprocessed by DrImpute, as supported by the significant increase of both POS and Kendall’s rank correlation score (Figure 3a). Figure 3b shows single cells of mouse early mesodermal development data on a 2D space using PCA without (left panel) and with (right panel) imputing the dropout events using DrImpute, and a pseudotime trajectory was constructed using TSCAN ([20]). Without imputation (left panel), the pseudotime trajectory started from E7.75 and ended at E7.75, which was not consistent with the known biological observations. On the other hand, with imputation (right), the pseudotime trajectory started from E6.5 and ended at E7.75, and both POS and Kendall’s rank correlation score significantly increased (POS increased from 0.66 to 0.89, and Kendall’s rank correlation increased from 0.5 to 0.63).

**Figure 3.**
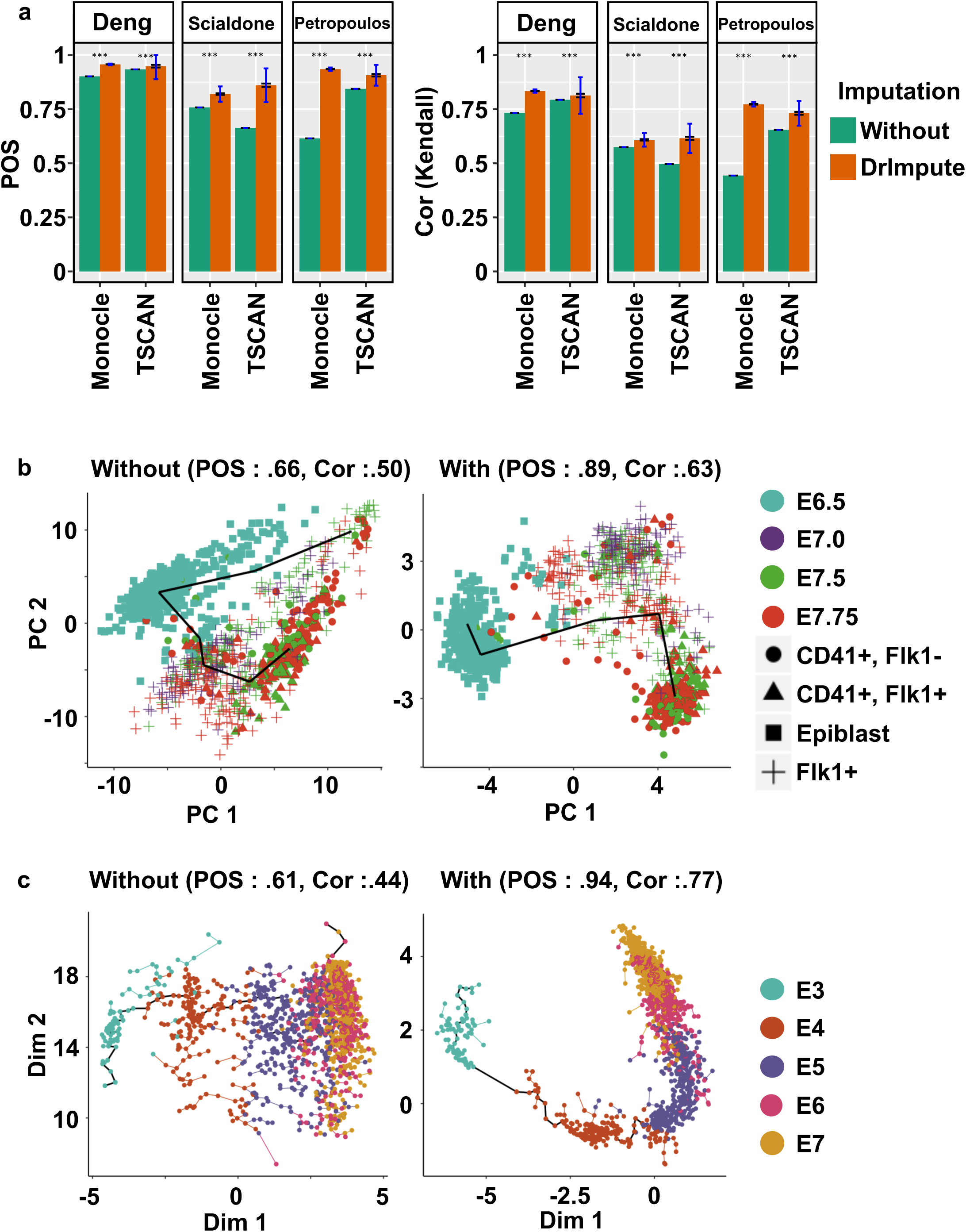
DrImpute greatly improved the performance of Monocle and TSCAN in lineage reconstruction. **(a)** The barplots of averaged Kendall’s rank correlation score and POS of 100 repeated runs of Monocle and TSCAN on three time series scRNA-seq datasets. Blue interval represents one plus or minus standard deviation of the category. Black interval represents one plus or minus standard error of the category. Both TSCAN and Monocle are deterministic with 0 variation before imputation. Wilcoxon rank sum test was utilized to compare Kendall’s rank correlation score and POS from different tools (⁎⁎⁎ *p* value < 0.001). (**b**) Visualization of lineage reconstruction of mouse early mesoderm ([52]) using TSCAN. The left and right panels, respectively, show the results of lineage reconstruction by TSCAN using the un-imputed scRNA-seq data or data preprocessed using DrImpute. **(c)** Visualization of lineage reconstruction for human preimplantation embryo ([53]) using Monocle. The left and right panels, respectively, show the results of lineage reconstruction by Monocle without and with preprocessing the scRNA-seq data using DrImpute.

As another example, Figure 3c showed single cells of human preimplantation embryo data on a 2D space using independent component analysis (ICA), with the pseudotime trajectory inferred by Monocle ([44]). When imputing dropout events using DrImpute (right panel), not only did the trajectory started from E3 and ended at E7, but the trajectory was also clearer in the sense that the E5, E6, and E7 stages were more separable compared to the trajectory inference results from non-imputed data (left panel). Consequently, the POS and Kendall’s rank correlation score were significantly increased (POS from 0.61 to 0.94; Kendall’s rank correlation from 0.44 to 0.77).

In summary, these results suggest that imputing dropout events using DrImpute also improved the performance of pseudotime inference using Monocle and TSCAN.

## Discussion

Dropout events and large cell-level variability are characteristic of scRNA-seq data, which are different from bulk RNA-seq data ([2]). However, many statistical tools derived for scRNA-seq data in cell type identification, visualization, and lineage reconstruction do not model for dropout events. Thus, we proposed a method for imputing dropout events considering cell-level correlation and systematically compared the performance without and with the imputation of dropout events. Our results on seven scRNA-seq datasets showed that imputing the dropout events using DrImpute significantly improved the performance of existing tools on cell type identification, visualization, and lineage reconstruction.

We would like to emphasize that DrImpute is the very first approach that sequentially utilizes dropout imputation with existing tools for more effective analysis. There are some statistical tools that model dropout events for specific purposes, such as BISCUIT, ZIFA, CIDR, and scImpute ([34], [35], [36], [18], [14], [37]). However, none of these suggest and compare the sequential use of dropout imputation and existing methods. We developed DrImpute to impute dropout events and demonstrated that the sequential use of dropout imputation employing DrImpute followed by the use of existing tools greatly improved the performance of the existing tools.

One of the limitations of DrImpute is that it considers only cell-level correlation using a simple hot deck approach. The gene-level correlation also exists, and more sophisticated modeling would improve the performance of the imputation. Most missing value imputation methods in bulk RNA-seq utilize gene-level correlation to impute missing values; for example, LLSimpute uses a local gene-level correlation structure to build local linear regression models to estimate missing values ([41]). One may improve the performance of DrImpute by modeling both cell-level and gene-level correlation.

## Conclusions

The main goal of the current study was to de-noise the biological noise in scRNA-seq data by imputing dropout events. We developed DrImpute and proposed the sequential use of DrImpute on existing tools that do not address dropout events. The results suggested that DrImpute greatly improved many existing statistical tools (pcaReduce, SC3, PCA, t-SNE, Monocle, and TSCAN) that do not address the dropout events in three popular research areas in scRNA-seq—cell clustering, visualization, and lineage reconstruction. In addition, DrImpute combined with pcaReduce, SC3 or t-SNE/kms showed higher performance in cell clustering than CIDR, which was specifically designed for the cell clustering of scRNA-seq data. DrImpute combined with PCA or t-SNE also demonstrated higher performance in 2D visualization than did ZIFA, which was specifically designed for the dimensional reduction of scRNA-seq data considering dropout events. Moreover, DrImpute imputed dropout events better than scImpute, as we have shown that the performance of t-SNE increased greatly in regard to cell clustering and visualization with DrImpute compared to that with scImpute. In summary, DrImpute can serve as a very useful addition to the currently existing statistical tools for single cell RNA-seq analysis.

## Methods

### Data preprocessing

Five scRNA-seq datasets (Pollen [10], Usoskin [39], Deng [6], Blakeley [7], and Treutlein [40]) were used for cell clustering and visualization, and three time series scRNA-seq datasets (Deng [6], Scialdone [47], and Petropoulos [48]) were used for lineage reconstruction. Genes that were expressed fewer than 2 cells were removed. The raw read counts were normalized by size factor[42], followed by log transformations (*log*_10_(*X* + 1)). Figure S2 shows data summaries, including how many zero entries are in the data, how many are imputed, and what is the computing time of DrImpute for each set of data.

### Imputation strategy

The general framework is described in Supplementary Figure 1, as follows: (1) computing the distance between cells using Spearman and Pearson correlations, (2) performing cell clustering based on a distance matrix, (3) imputing zero value cells multiple times based on multiple clustering results, and (4) averaging multiple imputation results to produce a final imputation for the dropout events.

Specifically, let *X* be an *n* by *p* log transformed gene expression matrix, where *n* is the number of rows (genes) and *p* is the number of columns (cells). The (*i*, *j*)th component of *X* is represented as *x*_*ij*_. Let *k* be the number of clustering results on data *X*, and *C*_1_, *C*_2_,…, *C*_*k*_ are each clustering results. Given that the clustering of *C*_*l*_ is a true hidden cell classification, the expected value of a dropout event can be obtained by averaging the entries in the given cell cluster:

*E*(*x*_*ij*_ *C*_*l*_ = *mean*(*x*_*ij*_|*x*_*ij*_ are in the same cell group in clustering *C*_*l*_). This step is also schematized in Supplementary Figure 1a. The *E*(*x*_*ij*_|*C*_*l*_) was computed for each clustering result *C*_1_, *C*_2_,…, *C*_*k*_, and the final imputation for the putative dropout events *x*_*ij*_, and *E*(*x*_*ij*_), was computed as a simple averaging:

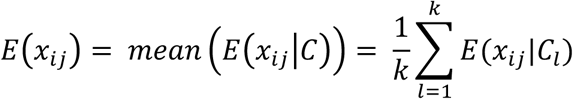

### Base clustering

For the default clustering of *C*_1_, *C*_2_,…, *C*_*k*_, we used an approach similar to that of SC3 ([11]). We first created a similarity matrix among cells using Pearson and Spearman correlations. K-means clustering was performed on the first 5% of the principal components of the similarity matrix and the number of clusters ranged from 10 to 15. Thus, the default setting had a total of 12 clustering results (two distance construction methods (Pearson, Spearman) times six numbers of clusters (10 to 15) for k-means clustering). This default setting was used for all the data analysis in this manuscript except for the down-sampling cells for the Blakeley dataset, which only had 30 cells. Its sample size was too low to use a default range for the number of clusters, so in this case, we used a clustering group size of 6 to 10.

### Software implementations and applications

The pcaReduce software was downloaded from the authors’ GitHub (<u>https://github.com/JustinaZ/pcaReduce</u>). We performed the analysis using the S and M options with the default setting. The SC3 package was downloaded from R Bioconductor (http://bioconductor.org/packages/release/bioc/html/SC3.html). To ensure the consistency of the comparison with other tools, the gene filtering option was turned off (gene.filter = FALSE). Other options were set as default. The Rtsne package and kmeans function in R program were used for t-SNE (perplexity = 9) followed by k-means. The log transformed expression data were centered as the gene level. We used the R kmeans function with the option iter.max = 1e+09 and nstart = 1000 for stable results. ZIFA software was then downloaded (<u>https://github.com/epierson9/ZIFA</u>). We used block_ZIFA with k = 15 for all data analysis, and we used the first two dimensions for visualization and evaluation. Monocle was downloaded from the R Bioconductor page (<u>https://bioconductor.org/packages/release/bioc/html/monocle.html</u>). In Monocle analysis, we first selected genes expressed in at least 50 cells and then selected differentially expressed genes using the differentialGeneTest() function (qval < 0.01). If there were no differentially expressed genes using the provided test, all genes expressing at least 50 cells were used for the subsequent analysis. TSCAN was downloaded from the R Bioconductor page (https://www.bioconductor.org/packages/release/bioc/html/TSCAN.html). All default settings were used for TSCAN.

### Evaluating the robustness of cell clustering

To evaluate the robustness of various cell clustering methods (Figure S3a and S3b), we down-sampled 100 genes (or about one-third of the cells; see Figure S2b for exact numbers) at random. PcaReduce, SC3, and t-SNE followed by k-means were applied to the down-sampled datasets, with or without imputing the dropout events using DrImpute and CIDR. The above processes were repeated for 100 times. The mean pairwise ARI of the clustering results from a total of 100 × 99 / 2 pairs of repeated runs was used as a robustness criterion using down-sampled genes (or cells). Note that when the cells were down-sampled, the overlapped cells were used for computing partial ARIs.

### Software availability

DrImpute was implemented as an R package, and it is available on CRAN (<u>http://cran.r-project.org)</u>. The core part of DrImpute is coded in C++ using the Rcpp ([43]) package.

## Supplementary Figure Legends

**Supplementary Figure 1.**
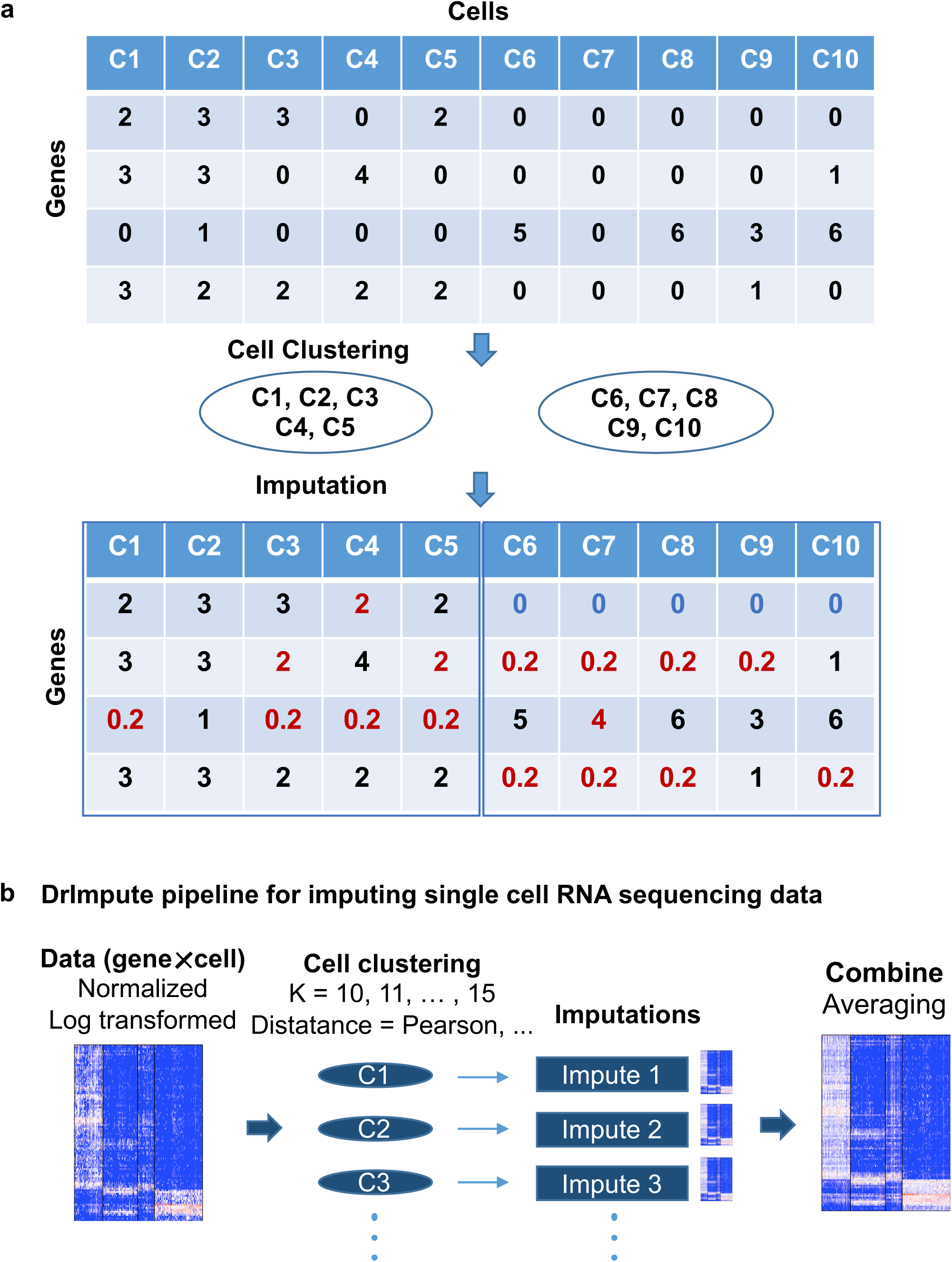
Basic procedure for clustering-based imputation. **(a)** Upper matrix is a gene by cell matrix. After clustering on gene by cell matrix, we get C1–C5 as one cluster and C6–C10 as the other cluster. Imputation is performed by averaging each cluster. **(b)** Overview of DrImpute pipeline: (1) perform data cleansing, normalizing, and log transformation; (2) calculate the distance matrix among cells; (3) impute the dropout entries based on the clustering results; and (4) average all imputation results to determine the final imputation.

**Supplementary Figure 2.**
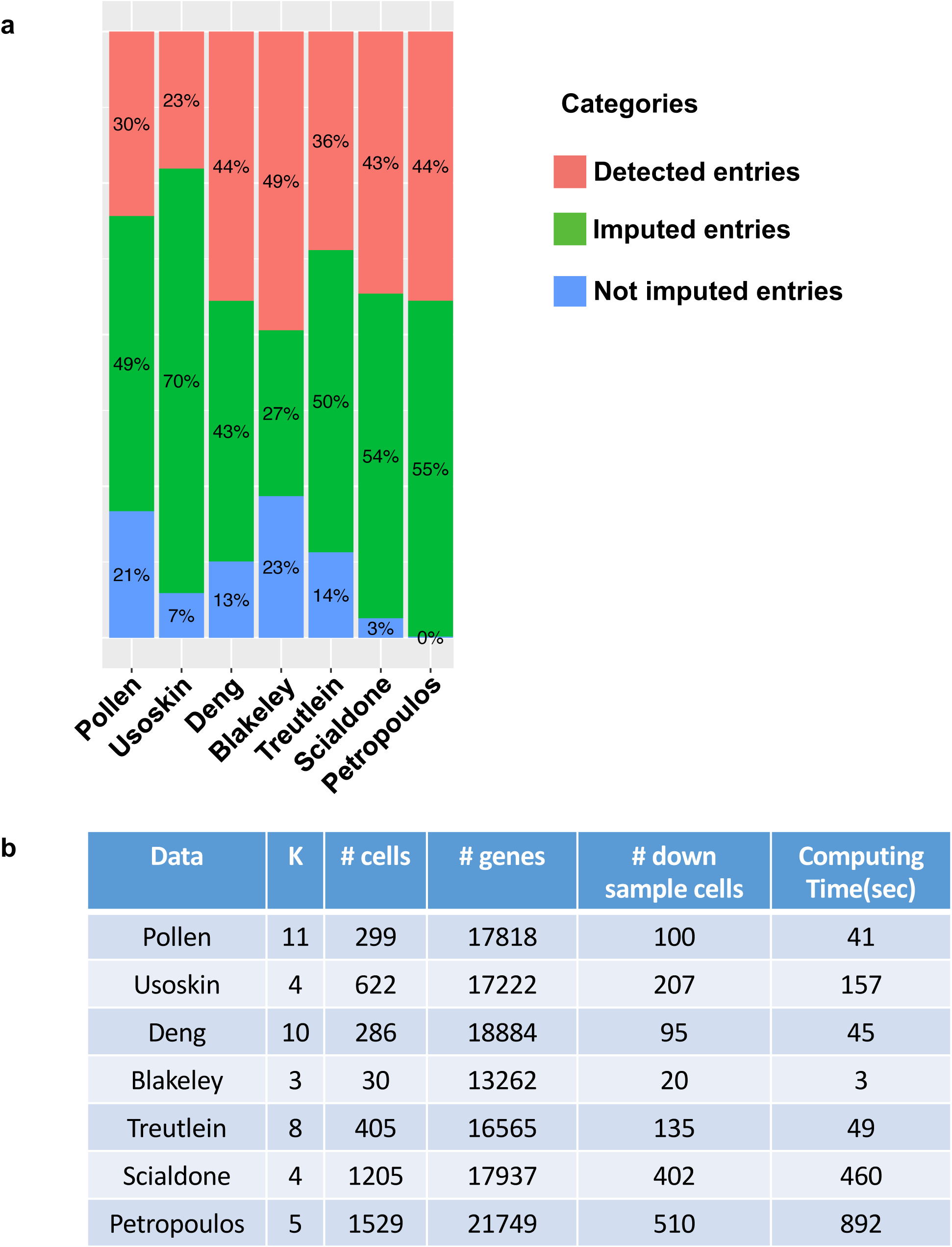
Overview of datasets used. **(a)** Percentage of detected, imputed, and not imputed entries for eight different scRNA-seq datasets. Genes that expressed less than 2 cells were previously removed. **(b)** The table describes the number of clusters, number of cells, number of genes, and number of down-sampled cells for robustness criteria and computing time of DrImpute for imputation. For the computing time, we used a single core CPU and averaged 100 runs using R microbenchmark package.

**Supplementary Figure 3.**
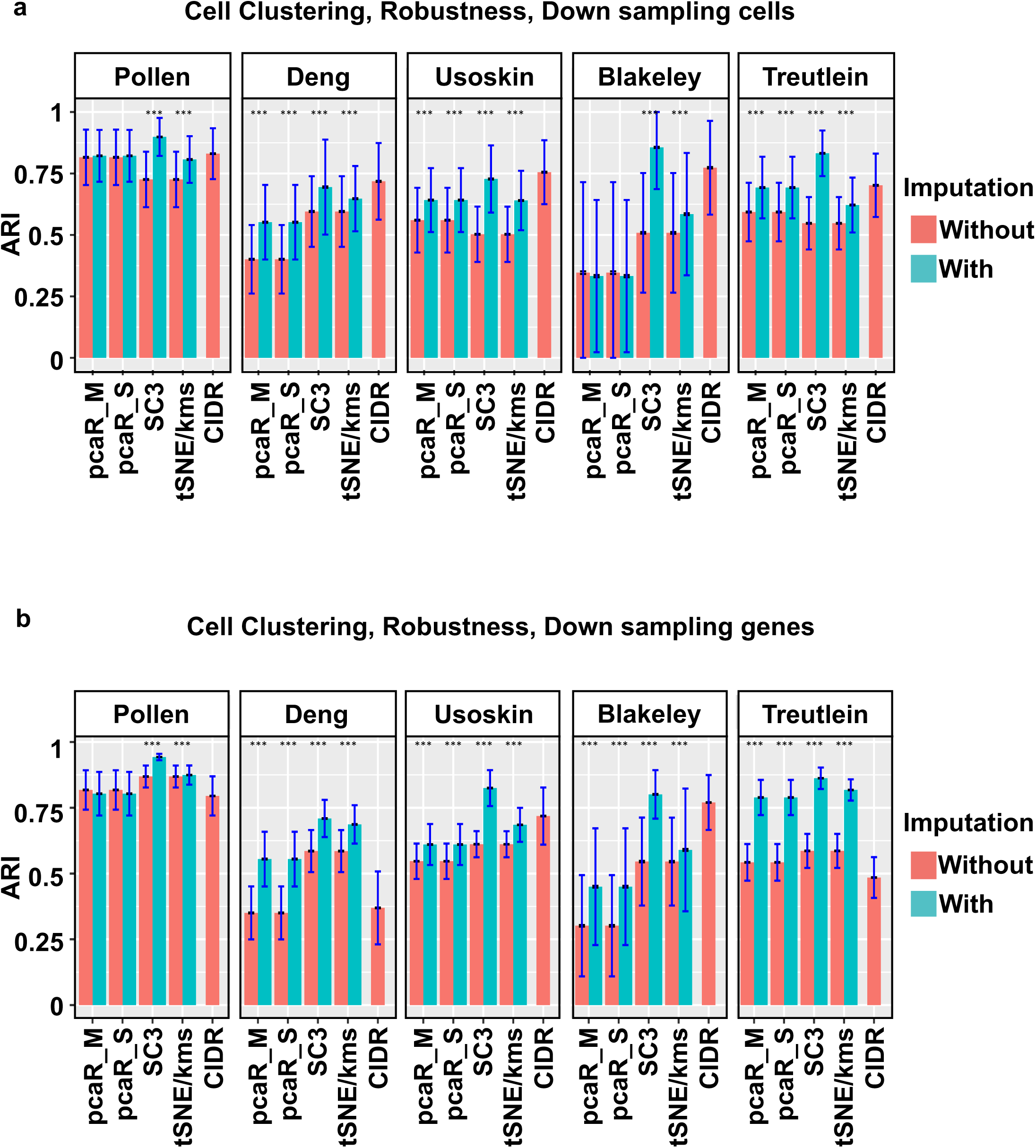
DrImpute significantly improved the performance of the existing tools for cell type identification in robustness criteria. To account for robustness, original datasets were down-sampled by cells (a) or by genes (b); we recorded clustering results for each data subset. ARIs are calculated for eah pair of data subsets. Barplot represents averaged ARIs. Blue interval represents one plus or minus standard deviation of the data. Black interval represents one plus or minus standard error of the data. Wilcoxon rank sum test is performed to compare before and after imputation. For down-sampled cells, 16 out of 20 cases are improved. For down-sampled genes, 18 out of 20 cases are improved (⁎⁎⁎ *p* value < 0.001).

